# Machine Learning Reveals Proteome-Encoded Growth Predictors of *Rhodopseudomonas palustris* CGA009 on Lignin Aromatics

**DOI:** 10.64898/2025.12.19.695510

**Authors:** Abraham Osinuga, Mark Kathol, Rajib Saha

## Abstract

Microbial utilization of lignin-derived aromatics requires extensive metabolic flexibility, yet growth outcomes vary sharply with substrate chemistry and oxygen availability. Whether this variability reflects distinct growth programs or alternative realizations of shared biochemical constraints remains unclear. Here, we combine quantitative proteomics with cross-condition machine learning to test whether growth-rate variation in *Rhodopseudomonas palustris* can be predicted directly from proteome composition and to identify the proteomic features that consistently encode growth potential across environments. Using OmniProt, a cross-condition neural modeling and interpretive framework, we predicted growth rates across 16 lignin-derived substrate–oxygen combinations, including held-out conditions, demonstrating that growth-relevant information is encoded in the proteome. Interpreting model reliance using Monte Carlo SHAP values and dependence-aware perturbation analyses revealed a compact, hierarchical organization of growth determinants. Despite pronounced oxygen-driven bifurcation in proteome abundance, the features required for accurate growth prediction were largely regime-invariant, defining a conserved biochemical core overlaid by adaptive, condition-specific modulators. These findings reconcile metabolic versatility with constrained growth control and show that diverse lignin-derived substrates converge onto a limited set of proteome-encoded growth bottlenecks accessed through flexible regulatory programs.

## Introduction

Lignin depolymerization releases a chemically heterogeneous pool of aromatic compounds that vary widely in redox state, energy yield, and catabolic accessibility (1, 2). *Rhodopseudomonas palustris*, a metabolically versatile phototrophic α-proteobacterium, is among the few bacteria capable of metabolizing many lignin-derived aromatics under both aerobic and anaerobic conditions (3–5). However, growth on these substrates is highly sensitive to oxygen availability, co-substrate support, and substrate chemistry, with closely related compounds supporting markedly different growth phenotypes. This variability suggests that growth limitation on lignin-derived substrates is governed not by pathway presence alone, but by higher-order constraints on metabolism, redox balance, and proteome allocation (6, 7).

Global transcriptomic and proteomic studies have revealed extensive induction and repression of metabolic pathways during growth on individual lignin-derived substrates. Yet such approaches often identify hundreds of responsive proteins without clarifying which changes are functionally informative for growth across environments (8, 9). Functional redundancy further complicates interpretation, as multiple transporters, oxidoreductases, and regulatory proteins can compensate for one another (10). Moreover, correlation with growth does not imply predictive or mechanistic relevance: many proteins track biomass accumulation or general metabolic demand rather than growth limitation per se. As a result, standard differential-expression and univariate analyses are poorly suited to identifying the subset of proteome features that remain predictively necessary across chemically and redox-diverse conditions (11). Similarly, metabolic (M) and metabolic expression (ME) models have been pivotal for advancing systems-level understanding in *R. palustris*—from characterizing substrate- and energy-dependent metabolic trade-offs using genome-scale metabolic reconstructions to revealing redox partitioning and electron allocation between carbon and nitrogen fixation pathways in the ME framework (9, 12). Yet, because these first-principles models fundamentally link metabolic flux to growth, they provide limited traction for evaluating how non-metabolic proteins—such as transporters, regulators, and stress-response factors—quantitatively influence growth across environments.

Here, we treat the proteome as an integrative readout of cellular state and ask whether growth-rate variation across lignin breakdown products can be predicted directly from proteomic composition—and, critically, which components of the proteome a generalizing model consistently relies upon to make those predictions. We introduce OmniProt, a cross-condition neural modeling and interpretive framework that couples cross-condition neural network prediction with dependence-aware feature attribution. OmniProt couples cross-condition neural network prediction with dependence-aware feature attribution to identify proteins whose quantitative variation encodes growth potential across environments. By focusing on predictive necessity rather than condition-specific responsiveness, this approach advances a systems-level view of metabolic versatility as emerging from flexible deployment of conserved growth-limiting processes.

## Results

### *R. palustris* grows on lignin breakdown products

We first quantified the capacity of *R. palustris* to grow on a panel of lignin breakdown products (LBPs) relevant to kraft lignin depolymerization. Cultures were supplied with p-coumarate (pC), p-coumaryl alcohol (pCA), coniferyl alcohol (CA), sinapyl alcohol (SA), sodium ferulate (SF), or kraft lignin (KL) as potential carbon sources, with acetate (Ac) included as a reference substrate due to its direct entry into central metabolism (13) (Fig. 1A). Growth curves were generated under aerobic and anaerobic conditions and fitted with logistic models (Supplementary Figure S1, Table S1), yielding condition-specific growth-rate parameters (Fig. 1A&C) that define the phenotypic space for subsequent predictive analyses. These measurements establish the energetic and physiological constraints imposed by chemically distinct lignin-derived substrates across redox regimes.

**Figure 1.**
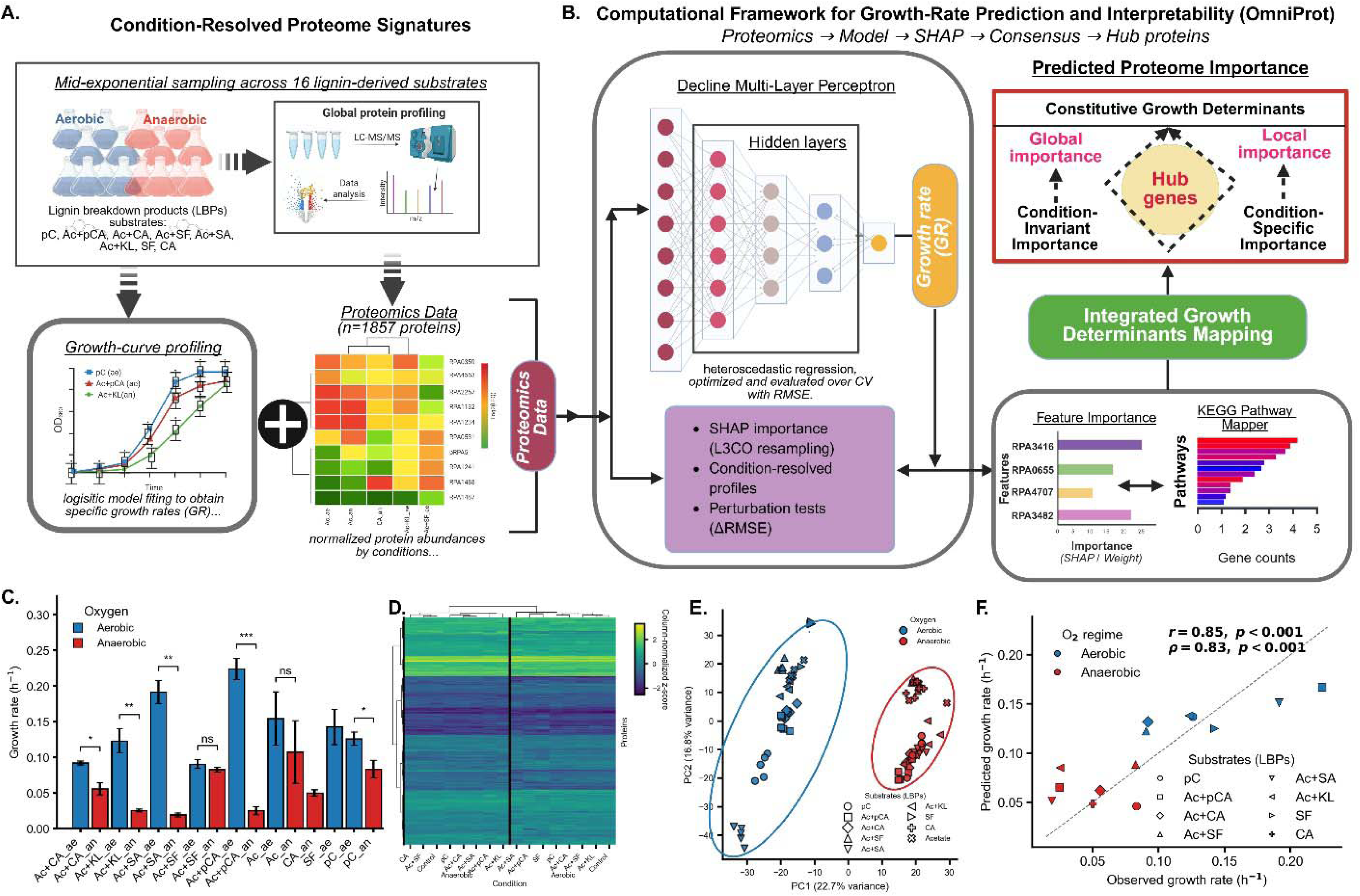
Experimental and computational framework of OmniProt. (A) R. *palustris* was grown across 16 lignin-derived substrates under aerobic or anaerobic conditions, yielding condition-resolved proteomes (1,857 proteins). (B) OmniProt trains a tapered MLP with condition-anchored cross-validation and Monte-Carlo SHAP to obtain global and condition-specific feature contributions, culminating in a refined hub set. (C–F) Growth-rate responses, proteome clustering, PCA separation of oxygen regimes, and high observed–predicted agreement demonstrate the biological structure captured by OmniProt. In (C) Statistical significance is denoted as p < 0.05 (*), p < 0.01 (**), p < 0.001 (***); ns, not significant.

Consistent with prior studies of lignin-derived aromatic utilization, several substrates failed to support sustained growth as sole carbon sources under either redox regime (4, 5). For these conditions, supplementation with acetate was required to supply ATP and reducing equivalents necessary to initiate peripheral aromatic catabolism. Importantly, acetate alone does not engage the aromatic degradation pathways encoded by *R. palustris*, as these genes are transcriptionally repressed or non-induced in the absence of an aromatic substrate, regardless of oxygen availability (14, 15). To ensure that lignin breakdown products contributed actively to biomass formation rather than serving as inert co-substrates, cultures supplemented with acetate were required to exceed the final optical density of acetate-only controls. Under this criterion, *p*-coumarate uniquely supported robust and reproducible growth without acetate under both aerobic and anaerobic conditions, whereas sodium ferulate and coniferyl alcohol supported more limited, condition-dependent growth in the absence of acetate, consistent with higher energetic or redox constraints on their assimilation (8, 10). In contrast, methoxylated monolignols and kraft lignin remained refractory, delineating substrate-specific energetic thresholds that constrain lignin-derived carbon assimilation in *R. palustris*.

### Proteome states associated with growth on lignin-derived substrates

Building on these growth phenotypes, we next characterized the proteomic configurations associated with lignin-derived substrate utilization. Five biological replicates were collected for each substrate–oxygen condition, with proteomic sampling performed at mid-exponential phase corresponding to the point of maximal instantaneous growth inferred from logistic fits (Fig. 1A, C). Proteomes collected at this stage capture the metabolic and regulatory state actively sustaining biomass accumulation under each condition while minimizing confounding effects of stationary-phase stress or substrate depletion (Fig. 1C-D). The resulting quantitative protein abundance profiles, comprising 1,857 proteins across 16 substrate–oxygen combinations, provide the feature space for all downstream modeling and interpretive analyses and enable direct alignment between substrate chemistry, growth physiology, and the proteomic states encoding condition-dependent metabolic adaptation.

### Oxygen availability reorganizes central metabolism and photosynthetic machinery

Because anaerobic growth in *R. palustris* requires engagement of a photoheterotrophic metabolic program, we next verified that cultures grown under anoxic conditions exhibited the expected physiological and proteomic signatures. Under anoxic conditions, we see strongly induced proteins associated with the photosynthetic reaction center and bacteriochlorophyll biosynthesis, including light-harvesting complex proteins *pucBA* (RPA2654) and *pucBC* (RPA3009), reaction-center subunits *pufL* (RPA1527) and *pufM* (RPA1528), bacteriochlorophyll biosynthesis enzymes *bchD* (RPA1507), *bchP* (RPA1532), *bchM* (RPA1546), and *bchE* (RPA1668), as well as carotenoid biosynthesis enzymes *crtI* (RPA1512), *crtB* (RPA1513), and *crtE* (RPA1519) —abundance patterns are shown in Supplementary Fig. S1c. Together, these components support light-driven electron flow and ATP generation via cyclic photophosphorylation. This transition was accompanied by reciprocal regulation of porphyrin biosynthesis enzymes, with repression of the oxygen-dependent coproporphyrinogen oxidase HemF (RPA1514) and induction of oxygen-independent isoforms HemN1 (RPA1666) and HemN2 (RPA0327), alongside reduced abundance of cytochrome *aa* oxidase components such as CoxB (RPA0831). Aromatic catabolism exhibited a similarly structured switch: aerobic conditions favored expression of β-ketoadipate pathway enzymes including *pcaF* (RPA0513) and *pcaC* (RPA4740), whereas anaerobic conditions selectively induced the benzoyl-CoA pathway regulator *badR* (RPA0655) and associated enzymes *badE* (RPA0658), *badD* (RPA0654), and *badI* (RPA0653). These findings indicate a strong activation of photosynthetic machinery and related pathways under anoxic, light-exposed conditions, consistent with *R. palustris* physiology. This remodeling was reflected globally in the proteome, with anaerobic conditions forming a coherent block in the clustered heatmap (Fig. 1C), separating cleanly from aerobic samples in PCA space (Fig. 1E), and across the full quantified proteome (Supplementary Fig. S3A), indicating a system-wide, oxygen-responsive reorganization of metabolism.

### Proteome composition quantitatively encodes growth potential across lignin substrates

To determine whether quantitative proteome states are sufficient to explain growth-rate variation across chemically diverse lignin breakdown products, we developed OmniProt (*Omni-conditional Proteomic modeling for growth trait inference*), a framework that links protein abundance profiles to physiological growth outcomes across environmental contexts. Using OmniProt, quantitative proteome profiles comprising 1,857 proteins were mapped to logistic growth-rate parameters measured across 16 substrate–oxygen combinations (Fig. 1B). Although *R. palustris* encodes ∼4,800 proteins, LC–MS/MS quantified fewer than 2,000 expressed proteins sufficient for growth prediction. Model performance was evaluated using an anchored leave-one-condition-out strategy in which all replicates from a held-out condition were excluded during training, while acetate reference conditions were retained in every fold to stabilize calibration across environments.

Across all folds, predicted growth rates closely tracked experimentally measured values (Fig. 1F), with a mean root mean squared error (RMSE) of 0.028 ± 0.018 h ¹. Given the observed growth-rate range of approximately 0.02–0.22 h ¹ across substrates and oxygen regimes (Fig. 1C), this error corresponds to ∼13–15% of the full physiological dynamic range and is comparable to the experimental variability observed within individual conditions. Importantly, this level of error is substantially smaller than the growth-rate differences separating aerobic and anaerobic states or distinguishing energetically favorable from constrained lignin substrates, indicating that the model resolves biologically meaningful variation rather than fitting noise. Condition-averaged predictions preserved both the absolute magnitudes and the rank ordering of growth outcomes across substrates and oxygen regimes (Pearson *r* = 0.85, *p* <0.001; Spearman ρ = 0.83, *p* = *p* <0.001). This agreement indicates that growth-rate differences across lignin-derived substrates are not dominated by idiosyncratic condition-specific effects but are instead encoded in structured, quantitative features of the proteome. As a result, this predictive sufficiency establishes a necessary foundation for mechanistic interpretation, implying that the subset of proteins repeatedly leveraged by the model represents constrained biochemical processes that actively limit growth across environments rather than passive correlates of substrate identity.

### Global determinant analysis separates regime-invariant growth constraints from adaptive predictors

Although accurate growth prediction establishes that growth-rate variation is encoded in the proteome, it does not by itself reveal which components of the proteome actively constrain growth across environments. Machine-learning models can assign high importance to proteins that merely track with particular substrates or oxygen regimes, conflating causal determinants with correlated passengers. To resolve this ambiguity, we developed a Global Determinant Analysis that evaluates persistence of model reliance across conditions (Fig. 1B; Fig. 2A). Importantly, “global” here refers to consistency in SHAP-derived (16) predictive contribution rather than invariance in protein abundance or direct biochemical causality.

**Figure 2.**
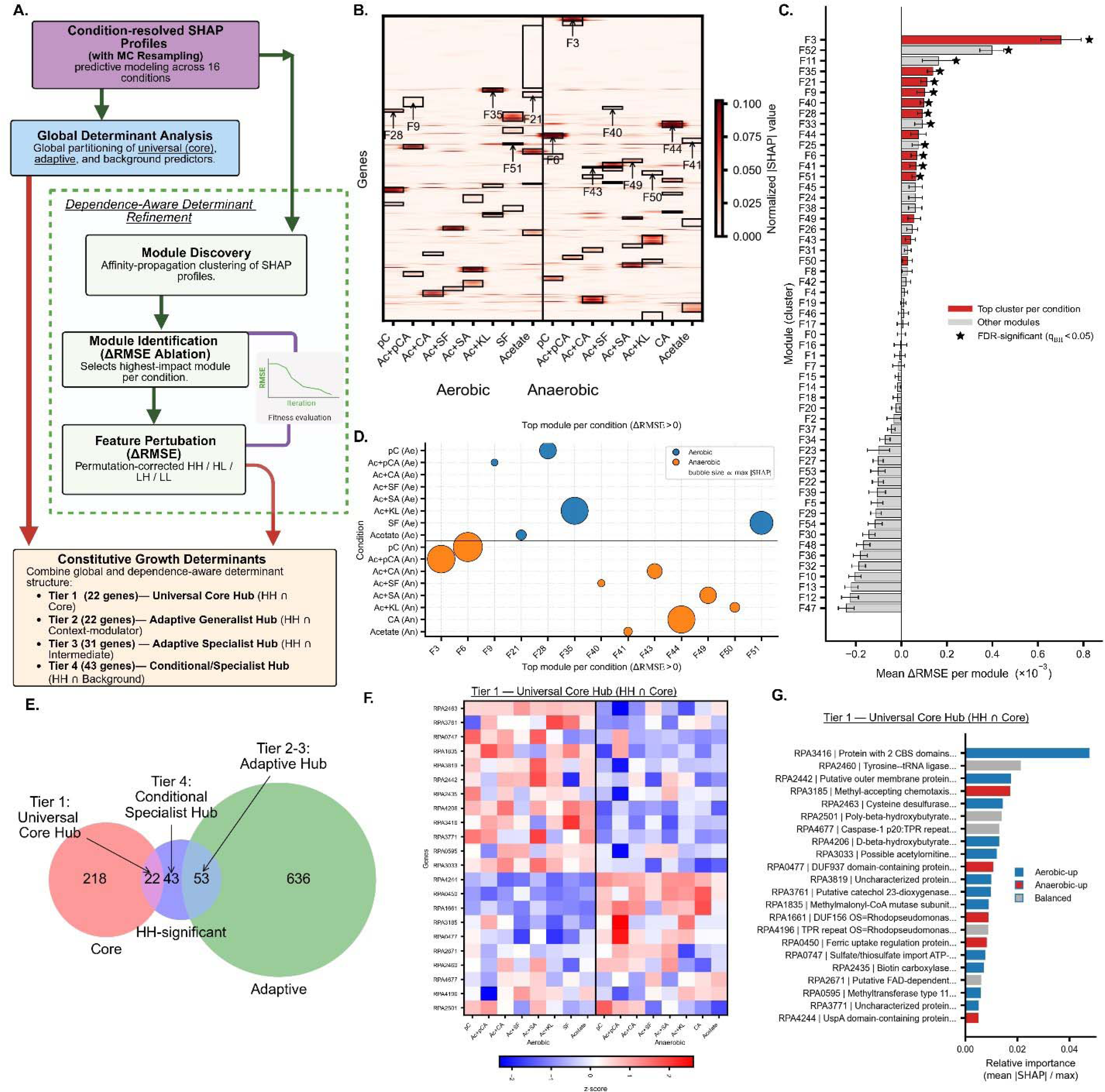
Modular and dependence-aware analysis identifies universal and context-linked growth determinants in *R. palustris*. **(A)** Integrated framework combining global determinant analysis with SHAP-derived module discovery, module ablation, and dependence-aware feature perturbation. **(B)** Condition-resolved SHAP profiles reveal modules with aerobic–anaerobic shifts in predictive influence. **(C)** Module ablation (ΔRMSE) identifies statistically significant predictive modules (FDR < 0.05). **(D)** Highest-impact module per condition highlights substrate-and oxygen-specific reliance patterns. **(E)** Four-tier hierarchy of Constitutive Growth Determinants: Tier 1 universal core, Tier 2 adaptive generalists, Tier 3 adaptive specialists, and Tier 4 conditional specialists. **(F)** Tier 1 Universal Core Hub: proteomic abundance profiles showing regime-invariant expression patterns, consistent with their universal predictive role. **(G)** Corresponding relative SHAP importance of the 22 high-confidence core members.

This approach identified a Universal Core Determinant set of 240 proteins exhibiting high global importance with minimal oxygen-associated variance, indicating that the model repeatedly relies on these features to constrain growth across conditions. Such regime-invariant reliance is consistent with proteome-allocation and bacterial growth-law frameworks, in which a restricted subset of biochemical processes imposes rate-limiting control across diverse physiological states (6, 7, 9, 17). In contrast, most proteins fell into background–intermediate predictors with diffuse influence (928 proteins) or adaptive modulators whose predictive contributions were moderate and shifted with substrate or oxygen context (225 and 464 proteins). Notably, after multiple-testing correction, no proteins exhibited predictive influence that was statistically exclusive to either aerobic or anaerobic conditions. This result reflects the structure of the *global determinant analysis*, which is defined in the space of SHAP-based feature reliance rather than protein abundance, and therefore does not contradict the strong oxygen-driven bifurcation observed in proteomic expression space (Fig. 1E). Instead, it highlights the unifying capacity of the model to identify shared growth-limiting constraints that persist across divergent expression programs.

### Growth control is concentrated in a limited number of non-redundant proteomic features

While the Global Determinant Analysis identifies proteins with persistent predictive influence, it does not establish whether that influence is statistically reliable or non-redundant at both condition and feature levels. Neural networks distribute importance across correlated predictors, and SHAP values alone cannot distinguish unique constraints from redundancy-mediated effects. To isolate proteins whose quantitative variation is truly required for prediction, we implemented a dependence-aware High-Confidence Determinant (HH) analysis combining module-level and feature-level conditional perturbation. This analysis first grouped proteins into coherent reliance patterns across environments using affinity propagation clustering of SHAP profiles, yielding 55 proteome-wide modules that capture distinct modes of model dependence (Fig. 2A–C). Module identity alone, however, does not imply functional necessity. We therefore systematically ablated each module and quantified the resulting loss of predictive performance, identifying a small subset whose removal produced statistically significant increases in error after false-discovery correction (Fig. 2C). Selecting the highest-impact module per substrate–oxygen condition revealed 13 predictive modules that collectively account for the model’s dependence across environments (Fig. 2D).

Within these predictive modules, we then resolved non-redundant determinants by applying a conditional perturbation framework that compares each protein’s effect under intra-module and full-proteome conditioning. This procedure isolates features whose predictive influence persists despite collinearity, pathway coupling, or shared regulatory structure. Applying these criteria identified 118 High-Confidence Determinants—a compact subset whose perturbation consistently degrades model performance across lignin substrates and oxygen regimes (BH-corrected q < 0.05; Fig. 2A). Fewer than 7% of quantified proteins satisfy this dependence-aware threshold, indicating that growth control in *R. palustris* is distributed across multiple processes but concentrated within a narrowly defined set of non-redundant constraints. While similar analyses in *E. coli* typically aggregate growth determinants into broad ‘sectors’ of the proteome (18), and gene signature studies in yeast often identify hundreds of correlated transcripts (19), our framework refines this to a compact ∼7% subset. This aligns with recent ‘sparse modularity’ findings in systems biology, which suggest that while thousands of genes respond to growth rate, actual control is distributed among a limited set of non-redundant bottlenecks (17). Importantly, the HH layer refines rather than recapitulates the Global Determinants: many globally influential proteins fail HH criteria due to redundancy, whereas several moderate contributors exhibit unexpectedly stable, non-substitutable influence. This separation establishes a rigorously defined substrate for mechanistic interpretation of growth-limiting processes beyond correlation structure alone.

### A hierarchical organization of growth determinants emerges from proteome-wide analysis

Despite the pronounced metabolic versatility of *Rhodopseudomonas palustris*, the oxygen-driven bifurcation of its proteome (Fig. 1E; Supplementary Fig. S3A) reflects a well-established role of oxygen as a primary organizer of bacterial proteome allocation and metabolic state (20, 21), but this separation alone does not identify which proteins actively constrain growth across environments. By integrating Global Determinant Analysis with the dependence-aware HH framework—both defined in the space of SHAP-derived model reliance rather than protein abundance—we resolved a compact, four-tier hierarchy of growth-relevant proteins that extends beyond correlative importance (Fig. 2A). The Universal Core Hub (Tier 1) is highly constrained, comprising only 22 high-confidence, regime-invariant proteins, in contrast to the 96 non-redundant Adaptive Modulators distributed across Tiers 2–4 (Fig. 2E). To place this hierarchy in functional context, all 118 HH-significant proteins were annotated against KEGG, classifying proteins as directly metabolic only when catalyzing registered reactions; consistent with the non-model status of *R. palustris* CGA009, many annotations remain putative. Only 42 proteins (36%) mapped to known metabolic pathways, whereas the majority (64%) comprised proteins of putative function, transcriptional regulation, or environmental sensing (Supplementary Table S4). This functional partition is reinforced by condition-resolved SHAP signatures (Fig. 2B–D), by the mixed oxygen-regime dependence defined by statistically significant differential abundance in the proteomic data (Fig. 2F-G), and by the persistence of structured—but attenuated—aerobic–anaerobic organization when analysis is restricted to the HH-significant subset (Supplementary Fig. S3B).

Across the four-tier hierarchy, mapped metabolic proteins are not uniformly distributed. Although the Universal Core Hub (Tier 1) contains a substantial fraction of metabolic enzymes (8 of 22; 36%), Tier 2 Adaptive Generalists Hub are comparatively enriched for non-metabolic or putative regulators (6 of 22 metabolic; 27%), consistent with broad, context-dependent modulation of core pathways rather than direct catalytic control. Tier 3 Adaptive Specialists (31 proteins) exhibit intermediate global importance with growth-predictive importance/influence confined to specific lignin substrates or oxygen regimes, reflecting localized enzymatic or regulatory functions, whereas Tier 4 Conditional/Specialist Hubs (43 proteins) display the highest metabolic density (17 of 43; 40%) yet contribute predictive power only under narrowly defined environmental conditions. This organization delineates a minimal, regime-invariant metabolic backbone that constrains growth across environments, overlaid by progressively specialized proteins that tune growth to substrate chemistry and redox state. Importantly, predictive influence remains distributed across a small set of high-leverage features rather than collapsing onto a single dominant protein (Supplementary Fig. S2), indicating that growth emerges from multiple partially redundant biochemical constraints rather than reliance on a single condition-linked proxy.

### A universal growth core links biosynthetic capacity, redox balance, and carbon assimilation

Integrating quantitative proteomic abundance patterns with the tiered hierarchy and condition-resolved model structure (Fig. 2B; Supplementary Fig. S3C–F) reveals growth control as a layered process rather than a diffuse property of the oxygen-partitioned proteome. Within the Universal Core Hub, several proteins maintain stable predictive influence across oxygen regimes while remaining non-redundant determinants of growth. These include tyrosyl-tRNA synthetase TyrS (RPA2460), reflecting sustained translational capacity, and poly-β-hydroxybutyrate synthase PhbC (RPA2501), which governs carbon storage and mobilization under fluctuating nutrient availability (22, 23). Other core members exhibit oxygen-associated abundance shifts yet retain regime-invariant predictive influence, indicating redistribution of conserved biochemical functions rather than substitution of growth constraints. These include cysteine desulfurase NifS1 (RPA2463), which supports Fe–S cluster assembly and thiamine biosynthesis (24); biotin carboxylase AccC (RPA2435), central to carboxylation reactions in fatty-acid and anaplerotic metabolism (25); methylmalonyl-CoA mutase MutB (RPA1835), which links propionate and branched-chain metabolism to central carbon flux (26); and iron- and stress-responsive regulators including Fur (RPA0450) and the UspA-like protein RPA4244, which coordinate iron homeostasis and stress adaptation under redox limitation (27, 28). The core further includes an FAD-dependent monooxygenase (RPA2671) whose abundance is invariant across oxygen regimes (Fig. 2F), linking cofactor biosynthesis to universal growth capacity, and a putative catechol 2,3-dioxygenase (RPA3761), which likely imposes a rate-limiting constraint on aerobic lignin-derived aromatic degradation across substrates, potentially enabling simultaneous engagement of meta- and ortho-cleavage routes to sustain high-flux aerobic degradation of chemically diverse lignin breakdown products. Collectively, these features define a conserved biochemical scaffold through which biosynthetic capacity, redox balance, and aromatic carbon assimilation jointly constrain growth, even as expression programs diverge across oxygen regimes.

### Adaptive regulatory and redox modules tune growth across substrate chemistry and oxygen availability

The second tier seems to resolve the regulatory interface through which oxygen availability reshapes growth control without redefining the core metabolic scaffold. Rather than imposing somewhat rate-limiting steps, proteins in this tier seem to modulate access to substrates, redox balance, and biosynthetic throughput within an otherwise conserved metabolic backbone. Accordingly, Tier 2 is enriched for transcriptional regulators, transport systems, and redox-balancing functions, and aligns closely with condition-specific SHAP growth predictive modules that dominate model reliance across environments (Fig. 2B; Fig. S2). Notably, the transcriptional regulator BadR (RPA0655) emerges as a prominent hub protein. BadR has been experimentally characterized as a regulator of anaerobic benzoate and aromatic acid metabolism in *R. palustris*, acting upstream of the benzoyl-CoA pathway and coordinating anaerobic aromatic catabolism under photoheterotrophic conditions (3, 8, 29–31). Additional regulators, including a GntR-family transcriptional regulator (RPA2343), alongside membrane and transport proteins such as an OprF-like porin (RPA4678) and a branched-chain amino-acid ABC transporter ATPase (RPA1791), indicate that growth modulation under oxygen limitation is governed primarily by regulatory gating of substrate access rather than large-scale replacement of core metabolic constraints (32). In parallel, this tier includes proteins associated with redox balance and biosynthetic throughput, including ferredoxin–NADP reductase Fpr (RPA1578), argininosuccinate synthase ArgG (RPA0392), acetolactate synthase IlvB (RPA3763), and ribosomal protein RpsD (RPA1589), consistent with established links between translational capacity, redox poise, and bacterial growth rate (18, 33, 34). The presence of 4-carboxymuconolactone decarboxylase PcaC (RPA4740) within this tier provides a direct biochemical anchor to oxygen-dependent β-ketoadipate pathway activity, a canonical route for aerobic aromatic funneling in diverse bacteria (35, 36). Collectively, these patterns indicate that aerobic adaptation recruits oxygen-enabled aromatic cleavage capacity, whereas anaerobic adaptation emphasizes regulated CoA-centered catabolism and redox/transport scaffolding—an interpretation consistent with established *R. palustris* physiology while highlighting a broad set of adaptive regulators elevated here as mechanistic candidates.

Moreover, the most substrate- and regime-specific biology is concentrated in the lower tiers 3 and 4, where specialized enzymatic and regulatory modules are recruited only under defined environmental demands. These proteins do not exert universal constraint on growth, but instead become necessary when particular metabolic routes, redox chemistries, or substrate entry points are engaged. Proteins associated with oxidative metabolism and central carbon flux—such as an NADH:ubiquinone oxidoreductase subunit (RPA2421), dihydrolipoyllysine succinyltransferase SucB (RPA0188), anthranilate synthase amidase TrpG (RPA4498), and a putative long-chain fatty-acid–CoA ligase (RPA2714)-are preferentially associated with oxygen-enabled metabolic states, consistent with established coupling between respiration, aromatic amino-acid biosynthesis, and growth (37–39). In contrast, proteins linked to iron acquisition (FbpA; RPA4152), protein maturation (peptide deformylase Def; RPA0621), chemotaxis (RPA4691), and redox-sensitive dehydrogenase activity (RPA1853) dominate under oxygen-limited conditions, reflecting strategies centered on resource acquisition and environmental sensing when oxidative chemistries are unavailable (40–42). The most localized, tier 4, further reveals lignin-aromatic entry and redox-buffering functions, including vanillate O-demethylase iron–sulfur subunit VanA (RPA3619), glutathione reductase Gor (RPA1983), glutathione S-transferase Gst2 (RPA0820), an acyl-CoA synthetase (RPA2302), and regulatory proteins such as GlnB (RPA2966) and CtrA (RPA1632). Vanillate O-demethylation via VanAB-type Rieske oxygenases represents a well-established entry point for methoxylated lignin-derived aromatics (43–45), whereas anaerobic aromatic metabolism in *R. palustris* proceeds through the benzoyl-CoA pathway under photoheterotrophic conditions (3, 29, 30, 35, 46). Many lignin-derived catabolic routes generate condition-specific carboxylic acid or CoA-ester intermediates (47), positioning RPA2302 as a versatile activator of these metabolites for downstream assimilation under anaerobic stress. Although no single tier reconstructs a complete lignin-degradation pathway, the hierarchy recovers recognizable oxygen-dependent aromatic funneling nodes and anaerobic regulatory architectures, while highlighting a substantial set of poorly annotated proteins whose context-specific recruitment represents a major and previously underexplored determinant of growth control.

### Conserved biochemical constraints underlie metabolic versatility

Despite the pronounced oxygen-driven bifurcation of the *Rhodopseudomonas palustris* proteome at the level of protein abundance (Fig. 1E), growth prediction reveals a strikingly different organization: the subset of features repeatedly required by the model to accurately predict growth remains largely conserved across environments. Aerobic and anaerobic conditions induce broad, system-wide remodeling of metabolic, photosynthetic, and respiratory proteins, yet these changes reorganize *how* metabolism is executed rather than *which* biochemical processes ultimately constrain growth. Importantly, this conclusion derives from analysis of model-based feature importance rather than differential protein abundance. The Global Determinant and High-Confidence analyses operate on SHAP profiles that quantify each protein’s contribution to prediction accuracy across conditions, clarifying that the absence of oxygen-exclusive global determinants reflects invariance in growth-limiting influence, not an absence of oxygen-dependent proteomic remodeling. This reconciliation demonstrates how distinct proteomic states can satisfy the same underlying growth constraints through alternative regulatory and metabolic configurations. By anchoring growth prediction to quantitative proteome features that remain informative across chemically and redox-diverse environments, OmniProt reveals growth as a structured outcome governed by a limited set of conserved biochemical bottlenecks, establishing a mechanistic foundation for interpreting metabolic versatility as flexibility layered atop invariant growth-limiting processes rather than wholesale rewiring of growth control.

## Discussion

This study demonstrates that metabolic versatility in *Rhodopseudomonas palustris* is not achieved through wholesale rewiring of growth control across environments, but rather through flexible regulatory and metabolic access to a conserved set of growth-limiting biochemical constraints. Although oxygen availability and substrate chemistry drive extensive remodeling of the proteome—most visibly reflected in the strong aerobic–anaerobic bifurcation of abundance space—the processes that ultimately limit growth remain largely invariant. By integrating quantitative proteomics with cross-condition learning and dependence-aware interpretability, OmniProt resolves this apparent contradiction and reveals growth as a structured, constrained outcome that is robust to environmental change, even as the underlying proteomic configuration varies substantially.

This perspective aligns with, and extends, classical theories of proteome allocation and bacterial growth laws, which posit that only a limited subset of cellular functions ultimately constrain growth rate across conditions. In *R. palustris*, these constraints span biosynthetic capacity, redox balance, translational throughput, and carbon assimilation, and are accessed through different regulatory realizations depending on oxygen availability and substrate chemistry. Aerobic and anaerobic states therefore represent alternative implementations of a shared growth architecture rather than distinct growth programs. Importantly, this interpretation emerges not from differential expression analysis alone, but from model-derived feature importance that identifies proteins whose quantitative variation the model repeatedly requires to resolve growth outcomes across environments. As a result, the conserved structure identified here reflects constraint at the level of growth prediction rather than uniformity of proteomic state. Notably, this work advances ANN-based methodological frameworks by replacing pooled correlational ranking with cross-condition generalization and dependence-aware hierarchy resolution of proteomic growth determinants (48).

A notable implication of this framework is that growth-relevant control is distributed across both well-characterized metabolic enzymes and a substantial number of poorly annotated or regulatory proteins, particularly within the adaptive and conditional tiers. While canonical aromatic catabolic pathways-such as oxygen-dependent β-ketoadipate funneling and anaerobic benzoyl-CoA metabolism-are clearly reflected in the hierarchy, no single tier reconstructs a complete lignin-degradation pathway. Instead, growth limitation appears to arise from flux-controlling entry points, redox-sensitive steps, and auxiliary functions that coordinate substrate activation, cofactor balance, and metabolic coupling. This finding underscores a key limitation of pathway-centric interpretations of metabolic versatility and highlights the importance of regulatory and accessory processes that are often underrepresented in genome-scale reconstructions, particularly in non-model organisms.

More broadly, the results suggest a shift in how metabolic adaptability should be interpreted and engineered. Rather than targeting individual pathway enzymes based solely on differential expression or pathway completeness, effective manipulation of growth and metabolism may require identifying and modulating the conserved biochemical bottlenecks that persist across environments, alongside the regulatory systems that gate access to them. By resolving growth control into invariant constraints and adaptive access mechanisms, OmniProt provides a generalizable framework for linking high-dimensional omics data to physiological outcomes in metabolically versatile organisms. This approach is readily extensible to other systems in which environmental plasticity masks conserved growth-limiting structure, offering a path toward more mechanistically grounded interpretation of data-driven models in systems biology.

## Limitations of the study

Several limitations of the present study warrant explicit consideration. First, SHAP-derived feature importance quantifies predictive necessity within the trained model and does not imply direct biochemical causality. Although the dependence-aware framework reduces the influence of correlated predictors, experimental perturbation will be required to establish causal growth limitation. Second, the proteomic measurements capture steady-state abundance at mid-exponential growth and do not resolve post-translational regulation, enzyme activity, or metabolite-level constraints that may further shape growth control. Third, pathway annotation for *R. palustris* CGA009 remains incomplete, and many proteins identified here as growth-relevant lack definitive functional assignment. Rather than weakening the conclusions, this gap emphasizes the value of model-guided prioritization for future biochemical characterization. Finally, the condition space explored here, while chemically diverse, is finite; extending this framework to additional substrates, stressors, or temporal dynamics will be necessary to assess the generality of the inferred constraints.

## Materials and Methods

### Growth experiments of *R. palustris* for different LBPs

*Rhodopseudomonas palustris* CGA009 (ATCC BAA-98) was used for all growth experiments. Strains were maintained at −80 °C in glycerol stocks (20% v/v for *R. palustris*) and recovered on 112 Van Niel’s medium, supplemented with appropriate antibiotics (49). For growth assays, aerobic seed cultures of *R. palustris* were established and subsequently inoculated into photosynthetic medium (The Conversion of Catechol and Protocatechuate to g-Ketoadipate by Pseudomonas putida) (PM) for lignin breakdown product (LBP) utilization experiments. Cultures were grown in 50 mL PM within 250 mL Erlenmeyer flasks under aerobic conditions or in 15 mL PM within sealed 16 mL balch tubes under anaerobic conditions. Anaerobic cultures were illuminated continuously using an LED shelf light (SN-AG230-WIR-065) in an Algaetron incubator (Photon System Instruments). All cultures were maintained at 30 °C with shaking at 275 rpm. PM was supplemented with 10 mM bicarbonate and 15.2 mM ammonium sulfate, and lignin-derived substrates were added at final concentrations of 1 mM as indicated. Where required, acetate (10 mM) was supplied to provide ATP and reducing equivalents necessary to initiate peripheral aromatic catabolism. To ensure that lignin breakdown products contributed directly to biomass formation rather than acting as inert co-substrates, cultures receiving both LBP and acetate were required to reach a final OD exceeding that of acetate-only controls (Supplementary Fig. S1). Growth curves represent the mean of three biological replicates.

### Proteome extraction, digestion, and LC–MS/MS

Cell pellets were lysed in Pierce RIPA buffer (Thermo Fisher Scientific) supplemented with 5 mM dithiothreitol and EDTA-free protease inhibitors (Roche) by heating at 95 °C for 10 min with agitation. Lysates were clarified by centrifugation (16,000 × g, 15 min), and protein concentrations were measured using the CB-X protein assay (G-Biosciences). Fifty micrograms of protein per sample were alkylated with 20 mM iodoacetamide (40 min, dark), quenched with dithiothreitol, precipitated with acetone, and washed three times with 70% ethanol. Pellets were resuspended in 50 mM Tris-HCl (pH 8.0) and digested sequentially with Lys-C (4 h) and trypsin overnight at 37 °C.

A pooled quality-control (QC) sample was generated by combining equal aliquots of all samples and injected after every 16 runs to monitor batch stability. Sample order was randomized by block randomization. Peptides were analyzed by nanoLC–MS/MS using an Ultimate 3000 RSLCnano system coupled to an Orbitrap Eclipse mass spectrometer (Thermo Fisher Scientific). Peptides were trapped on an Acclaim PepMap 100 column (75 µm × 2 cm) and separated on a C18 nano column (Peptide CSH, 75 µm × 250 mm) at 300 nL min ¹ using a 75-min gradient from 5% to 22% acetonitrile in 0.1% formic acid. Data were acquired in data-dependent mode with MS¹ scans at 120,000 resolution (m/z 375–1500) followed by HCD MS² acquisition in the ion trap.

### Protein identification and preprocessing

Protein identification and quantification were performed using Proteome Discoverer v2.4 with Mascot against a combined *R. palustris* CGA009 UniProt database (UP000001426_258594) and a modified cRAP contaminant database. Searches assumed trypsin digestion with up to two missed cleavages, precursor tolerance of 15 ppm, and fragment tolerance of 0.06 Da. Carbamidomethylation of cysteine was set as a fixed modification, with methionine oxidation and asparagine/glutamine deamidation as variable modifications. Peptide-spectrum matches were filtered to 1% false discovery rate using Percolator. Only proteins identified by at least two unique peptides and five peptide-spectrum matches were retained.

Quantitative proteomics data were processed in MaxQuant Perseus. Proteins were required to be present in at least three replicates across all sample groups; others were discarded. Label-free quantification (LFQ) protein intensities (our proxy abundances here) were log -transformed and missing values imputed using a downshift of 1.8 standard deviations and a width of 0.3 standard deviations. Quantile normalization was performed in R. Differential abundance between aerobic and anaerobic conditions was assessed on globally z-scored abundances using Welch’s two-sample t-test with Benjamini–Hochberg correction (FDR < 0.05), classifying proteins as aerobically or anaerobically enriched based on the sign of the mean difference.

### Model architecture, training, and cross-condition generalization

Proteomic data were analyzed using OmniProt, a learning and interpretive framework that maps condition-resolved proteome states to quantitative growth phenotypes while identifying features required for cross-environment generalization. The final dataset comprised 80 proteomic profiles spanning 16 substrate–oxygen conditions, quantified across 1,857 proteins. Logistic growth-rate means and standard deviations were used as response variables, and condition metadata were retained to enforce condition-aware data partitioning.

Growth prediction employed a tapered multilayer perceptron (Decline-MLP) with three fully connected layers (311→120→46 units), ReLU activation, dropout (p = 0.1186), and the AdamW optimizer (learning rate 2.34 × 10 ; weight decay 1.47 × 10 ). Models were trained for up to 1,000 epochs with early stopping based on internal validation loss. A heteroscedastic Gaussian negative log-likelihood loss was used to account for replicate-level uncertainty in measured growth rates, jointly predicting mean μ(x) and variance σ²(x):

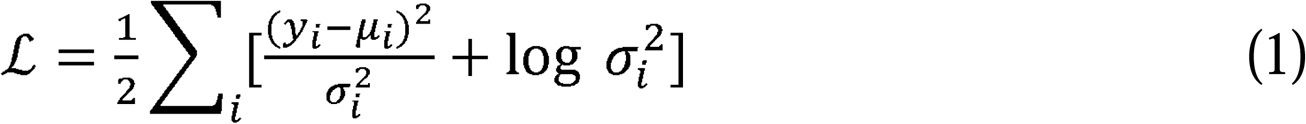

Generalization was evaluated using exhaustive leave-one-condition-out cross-validation (LOOCV). Each of 14 environmental conditions was held out in turn, excluding all its replicates from training. Remaining conditions were split into internal training (∼85%) and validation (∼15%) sets. Two acetate reference conditions (*Ac_ae* and *Ac_an*) were included in every training fold to stabilize calibration and prevent fold-to-fold drift. Performance metrics (RMSE, Pearson and Spearman correlations) were computed at both sample and condition-mean levels and summarized across folds. Model hyperparameters (learning rate, layer decay, dropout, weight decay, and initial hidden width) were optimized using Optuna with a TPE search (random seed = 42). Proteomic features were standardized to zero mean and unit variance prior to training.

### Monte Carlo SHAP analysis and growth-determinant analyses

Condition-resolved protein contributions to growth prediction were quantified using a Monte Carlo SHAP (SHapley Additive exPlanations) framework (see Supplementary Methods S1). Briefly, models were repeatedly retrained under leave-three-conditions-out cross-validation, in which all biological replicates from three non-anchor substrate–oxygen conditions were excluded per iteration, while acetate reference conditions (*Ac_ae*, *Ac_an*) were retained to stabilize calibration. Kernel SHAP was applied to held-out samples using training data as the background distribution, producing condition-resolved contribution profiles for all quantified proteins. These SHAP-derived importance profiles, rather than protein abundances, formed the basis for all downstream analyses.

Global growth determinants were identified by aggregating condition-resolved SHAP profiles across all environments and decomposing importance variance into regime-associated and condition-specific components (Supplementary Methods S2). Proteins were classified into aerobic, anaerobic, oxygen-bridging, or adaptive determinant classes based on global importance, regime variance, and aerobic–anaerobic effect size. High-Confidence Determinant (HH) analysis further refined this set using SHAP-based module construction and dependence-aware conditional perturbation to identify proteins whose quantitative variation was non-redundantly required for accurate growth prediction across environments (Supplementary Methods S3–S6).

### Simulation platform

All computational analyses for the **OmniProt** pipeline were conducted in Python (v3.10.18) using PyTorch (v2.4.1) with CUDA 11.8 and cuDNN 9.1, executed on the University of Nebraska–Lincoln’s Holland Computing Center (HCC) SWAN cluster running Ubuntu Linux. Core models were trained on an NVIDIA Tesla V100S GPU (32 GB). The computational node contained dual Intel(R) Xeon(R) Gold 6248R CPUs (20 cores allocated per job) and 64 GiB of RAM, providing sufficient throughput for large-scale cross-condition learning, anchored LOOCV procedures, and SHAP-based interpretability analyses. This standardized environment enabled consistent benchmarking and reproducible execution of all OmniProt experiments.

## Data Availability

All the codes required to reproduce the results of the manuscript can be accesses at https://github.com/ssbio/OmniProt

## Acknowledgements

This work was completed utilizing the Holland Computing Center of the University of Nebraska, which receives support from the UNL Office of Research and Innovation, and the Nebraska Research Initiative. We gratefully acknowledge funding support from the National Science Foundation (NSF) CAREER grant 1943310 and NIH MIRA Award (5R35GM143009) both awarded to R.S.

---

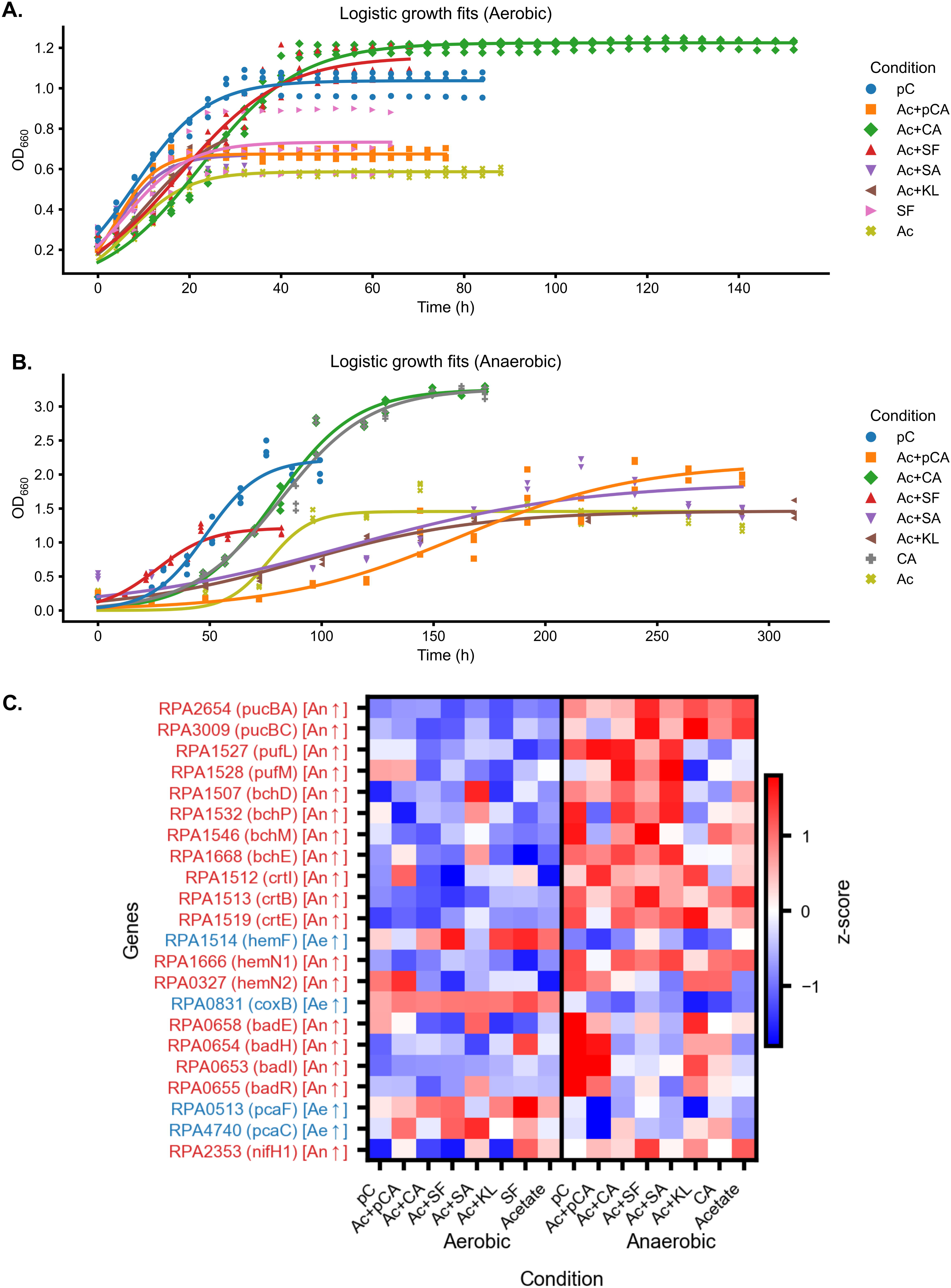

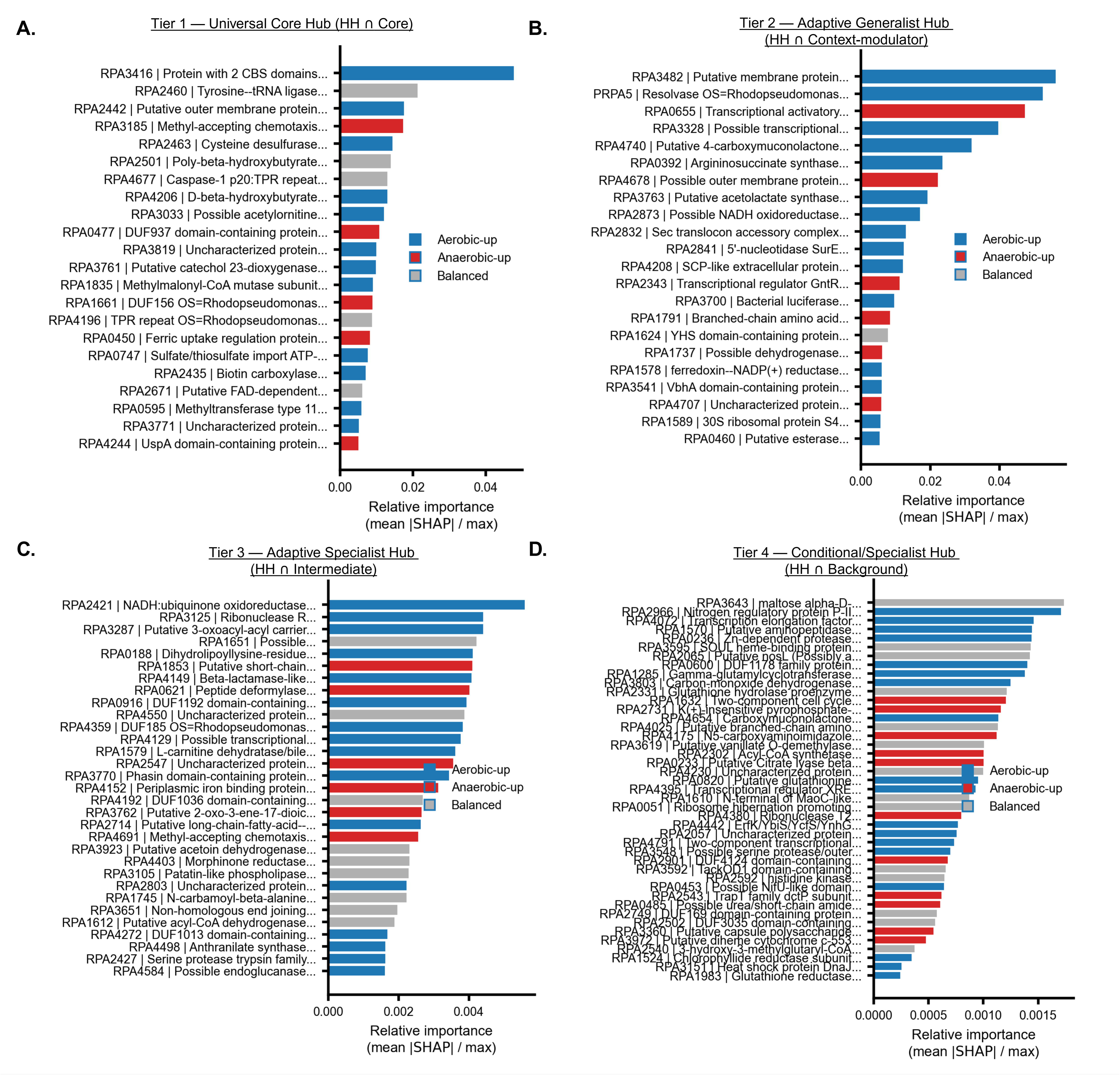

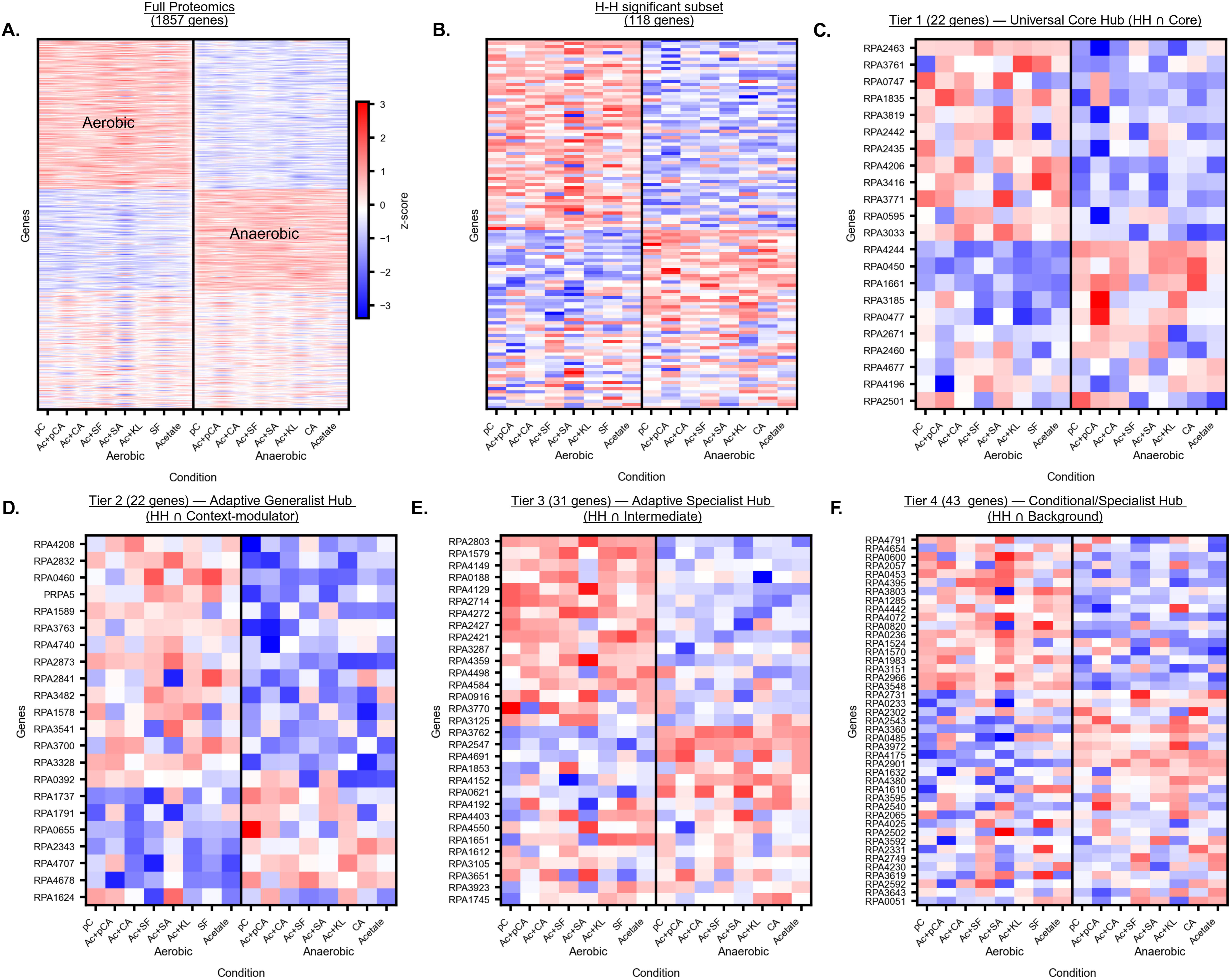

