## Supporting Information for "Machine Learning Reveals Proteome-Encoded Growth Predictors of *Rhodopseudomonas palustris* CGA009 on Lignin Aromatics"

Running Title: Proteome-informed growth modeling in *R. palustris* CGA009

This Supplementary Information supports the manuscript:

**Supplementary Figures**

- **Figures S1 – S3**

**Supplementary Methods**

- **S1**. Monte-Carlo SHAP analysis workflow
- **S2.** Global Determinant Analysis framework
- **S3.** High-Confidence Determinant (HH) analysis
- **S4.** SHAP-based module construction
- **S5.** Conditional perturbation testing workflow
- **S6.** Feature-level redundancy classification

**Supplementary Tables**

Tables **S1–S12** include: growth measurements, quantitative proteomics, module assignments, determinant tiers, and pathway annotations across all substrate–oxygen conditions.

### Supporting Information

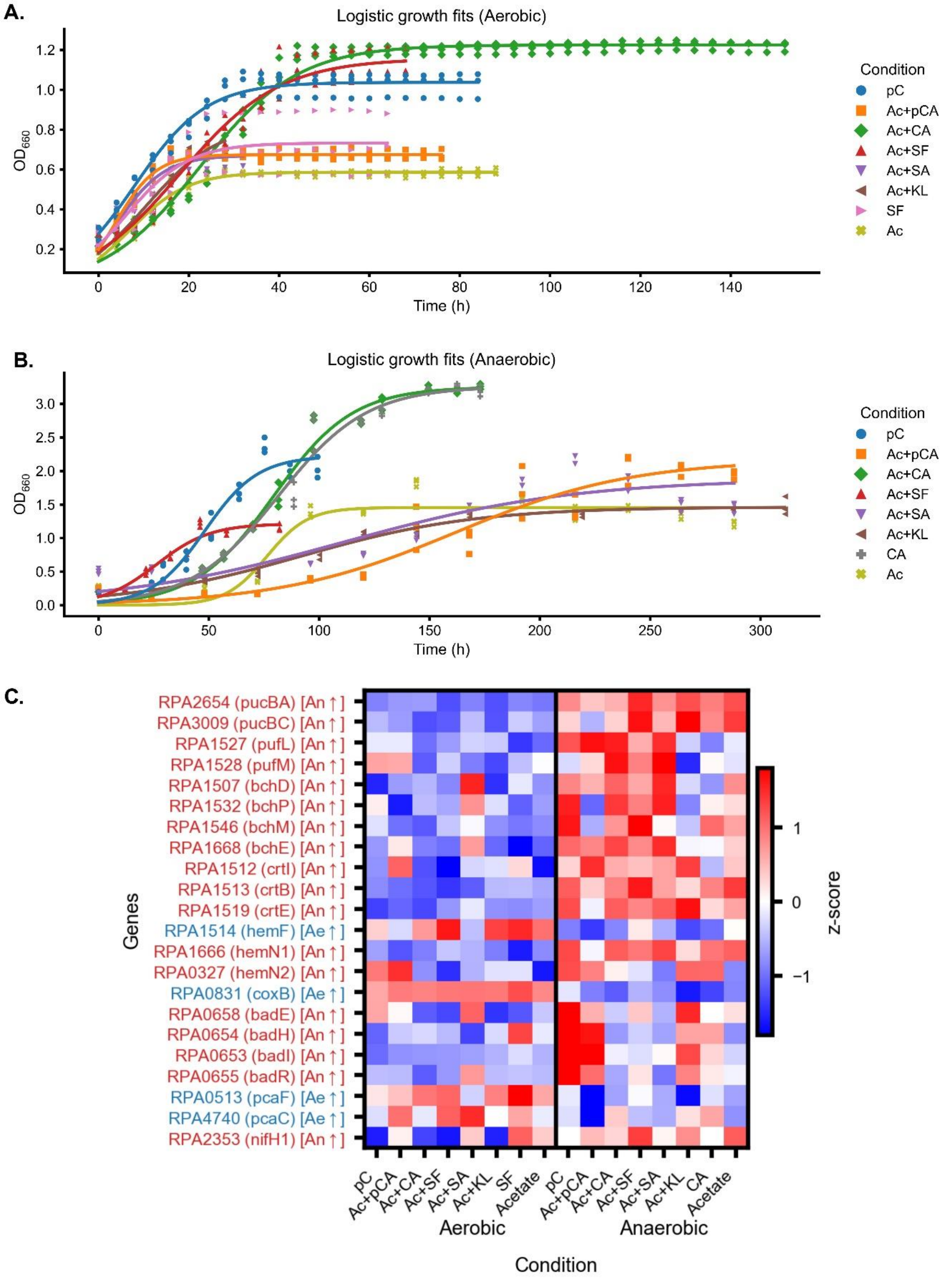


Figure S1. **Growth phenotypes and oxygen-dependent proteomic signatures during lignin breakdown product utilization in *Rhodopseudomonas palustris*. (A)** Logistic growth fits for aerobic cultures grown on lignin-derived substrates with or without acetate; points denote measured $OD_{660}$ values and lines indicate fitted models. **(B)** Corresponding growth dynamics under anaerobic, light-exposed conditions, revealing distinct substrate- and oxygen-dependent growth behaviors. **(C)** Heatmap of representative proteins showing oxygen-dependent proteomic reprogramming, including photosynthetic apparatus, bacteriochlorophyll biosynthesis, porphyrin metabolism, and aromatic catabolic pathways. Protein abundances are shown as row-wise z-scores, grouped by oxygen regime and substrate condition.

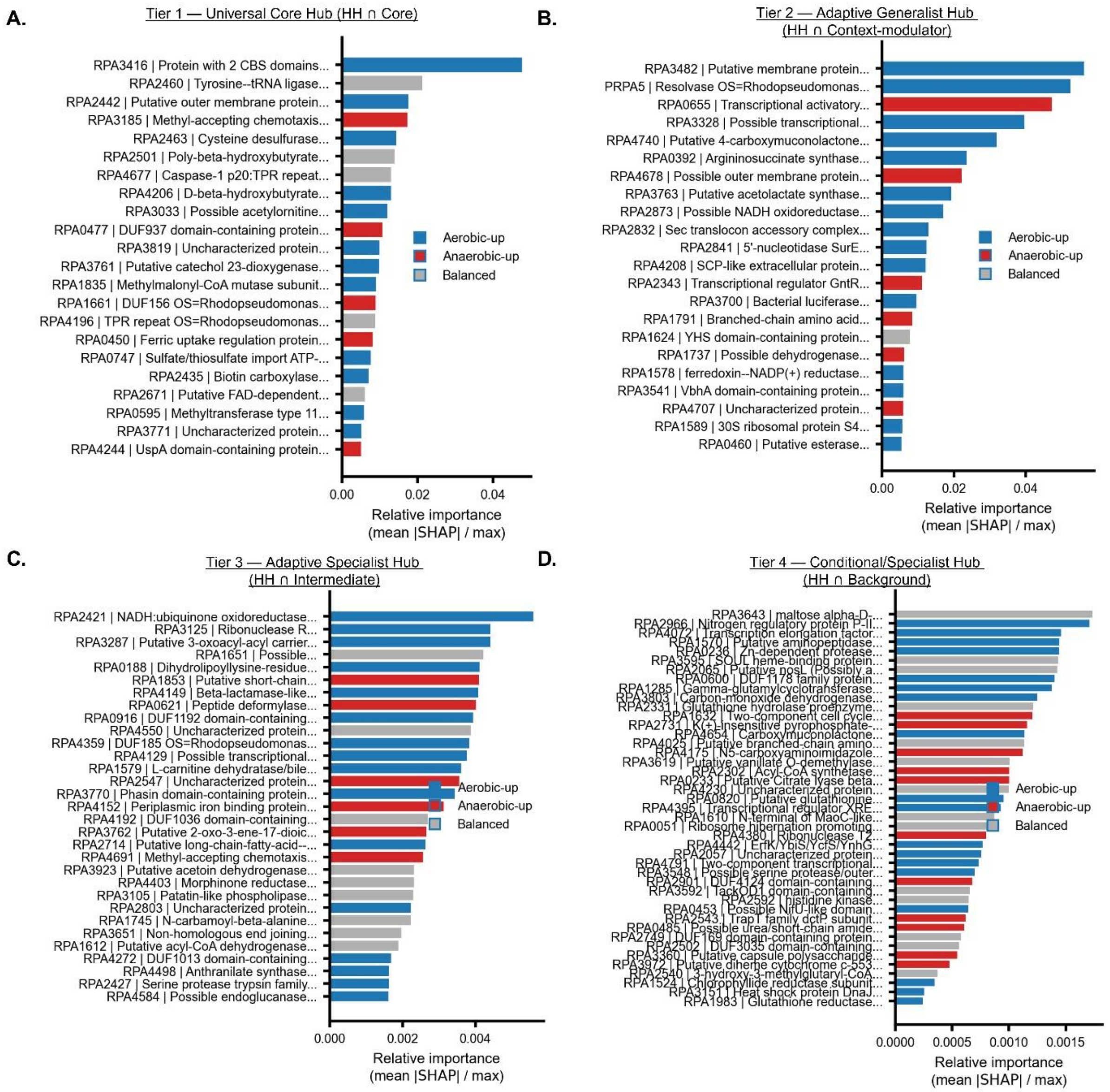


Figure S2. **Hierarchical organization of growth determinants identified by dependence-aware model interpretability. (A–D)** Ranked protein features within the four determinant tiers: **Tier 1**, Universal Core Hub; **Tier 2**, Adaptive Generalist Hub; **Tier 3**, Adaptive Specialist Hub; and **Tier 4**, Conditional/Specialist Hub. Bars indicate relative predictive importance (mean |SHAP| normalized to the global maximum). Colors indicate statistically significant differential protein abundance between oxygen regimes (aerobic-up, anaerobic-up; FDR < 0.05), or no significant difference (balanced).

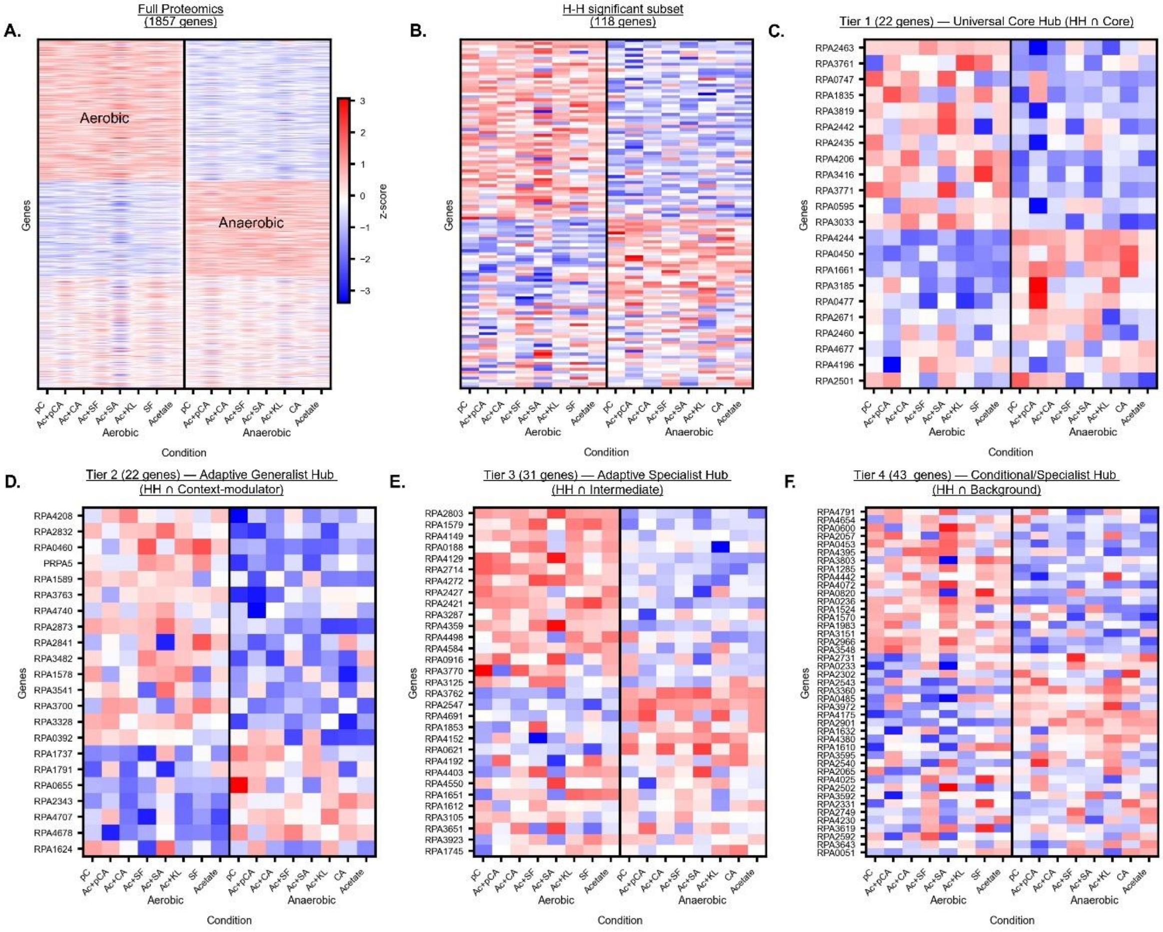


Figure S3. **Proteome-wide oxygen structuring and hierarchical refinement of growth determinants. (A)** Full proteome (1,857 proteins) shows strong oxygen-driven bifurcation in abundance space. **(B)** The HH-significant subset (118 proteins) retains oxygen structure while substantially reducing dimensionality. **(C–F)** Condition-resolved abundance patterns for proteins in the four determinant tiers: **Tier 1**, Universal Core Hub (22 proteins); **Tier 2**, Adaptive Generalist Hub (22 proteins); **Tier 3**, Adaptive Specialist Hub (31 proteins); and **Tier 4**, Conditional/Specialist Hub (43 proteins). Colors indicate z-scored protein abundance across substrate–oxygen conditions. These panels illustrate how strong oxygen-dependent proteomic remodeling coexists with a compact, hierarchically organized set of growth-relevant features identified through dependence-aware model interpretation.

**Supplementary Tables S1–S12** provide all supporting datasets for this study, including growth measurements, quantitative proteomics, model-derived determinant tiers, and pathway annotations for *Rhodopseudomonas palustris* CGA009 across lignin-derived substrates and oxygen regimes.

#### Supplementary Methods S1 — Monte Carlo SHAP analysis for condition-resolved contribution profiles

To quantify condition-resolved protein contributions, Monte Carlo SHAP analysis was performed. All non-anchor conditions were enumerated, and every unique combination of three conditions was held out, yielding 364 leave-three-conditions-out splits. For each split, the Decline-MLP was retrained de novo, retaining the acetate anchors. Kernel SHAP was applied to held-out samples using training samples as the background distribution, with 5,460 model evaluations per instance. SHAP values were computed for all proteins and aggregated across splits to obtain global and condition-specific contribution profiles. These SHAP-derived profiles, rather than protein abundances, were used for all downstream analyses.

To quantify how proteome composition contributes to growth-rate prediction across environmental contexts, we performed a Monte Carlo SHAP (SHapley Additive exPlanations) analysis prior to any clustering or module construction. SHAP decomposes model predictions into additive, sample-specific feature contributions, providing a principled measure of how individual proteins influence predicted growth rates independent of their absolute abundance. To rigorously probe extrapolative behavior, all non-anchor substrate–oxygen conditions were enumerated, and every unique combination of three conditions was treated as a held-out group, yielding 364 leave-three-conditions-out (L3CO) splits. In each split, all biological replicates from the selected conditions were excluded entirely from model fitting, while the two acetate reference conditions (Ac_ae and Ac_an) were retained in every training fold to stabilize calibration. The remaining conditions were partitioned at the condition level into internal training and validation sets, typically comprising eight training and three validation conditions per iteration, ensuring that no sample from a held-out condition influenced parameter estimation or early stopping.

For each L3CO split, the Decline-MLP was reinitialized and trained de novo using the same architecture, optimizer, and heteroscedastic loss formulation described above. Kernel SHAP was then applied to samples from the held-out conditions, using all samples from the corresponding training fold as the background distribution and test samples as instances to be explained. Each instance was evaluated under a fixed computational budget of 5,460 model calls, and SHAP values were computed for all 1,857 quantified proteins. This procedure yielded high-resolution, condition-aware estimates of how variation in individual protein abundances contributes to predicted growth rates, as learned by independently trained models.

Global contribution profiles were obtained by aggregating absolute SHAP values across all explained samples and Monte Carlo iterations and normalizing by the total number of evaluated instances. To preserve biological context, SHAP values were also averaged within each substrate–oxygen condition, producing a condition-by-protein matrix of mean absolute contributions. These condition-resolved SHAP profiles formed the foundation for all downstream analyses, including identification of regime-invariant versus context-dependent predictors, module discovery, and dependence-aware perturbation tests. Importantly, because these analyses operate on model-derived contribution profiles rather than on differential protein abundance, they isolate features that are repeatedly required for accurate growth prediction across environments rather than those that merely co-vary with specific conditions.

#### Supplementary Methods S2 — Global Determinant Analysis: pan-condition importance metrics and regime-variance decomposition

To identify **global growth determinants**—proteins whose predictive influence persists across substrates and oxygen regimes—we consolidated condition-resolved SHAP perturbation results into a set of pan-condition importance metrics. Aggregating SHAP profiles across all 16 substrate–oxygen conditions yielded a condition-by-protein matrix in which each entry reflects the magnitude of the model's reliance on a given protein within a specific environmental context. Because SHAP values are derived from independently trained models and therefore differ in scale, variance structure, and correlation geometry, they are not directly comparable across conditions. We therefore implemented an ANOVA-style variance decomposition framework to define global determinants based on model-derived importance structure rather than protein abundance.

Condition-wise SHAP magnitudes were first normalized by their respective median absolute values to enable comparability across heterogeneous substrates and oxygen regimes. From this normalized matrix, we computed for each protein (i) a global mean importance across all conditions, (ii) aerobic- and anaerobic-specific mean importance values, and (iii) their difference (ΔAn–Ae). Total importance variance was then partitioned into between-regime and within-regime components, yielding a regime variance fraction that quantifies the extent to which a protein's predictive relevance is explained by oxygen availability rather than substrate-specific or idiosyncratic effects. Statistical significance of regime-associated biases was assessed using Welch's t-tests comparing aerobic and anaerobic SHAP distributions for each protein, with multiple testing controlled by the Benjamini–Hochberg procedure (FDR $< 0.05$).

Proteins were subsequently classified using data-driven percentile thresholds applied to global mean importance, regime variance fraction, and aerobic–anaerobic effect size. High-importance proteins were defined as those exceeding the 75th percentile of global mean importance, whereas proteins below the median were designated background contributors. Among high-importance proteins, those exhibiting significant regime bias ($q < 0.05$) and high regime variance (top quartile) were classified as aerobic or anaerobic global determinants, depending on the sign of ΔAn–Ae. High-importance proteins with non-significant regime effects and low regime variance were classified as oxygen-bridging pan-condition global determinants, while remaining high-importance proteins were designated adaptive global modulators. Importantly, this Global Determinant Analysis operates entirely on SHAP-derived importance profiles and does not imply regime-invariant protein abundance, but rather identifies proteins whose predictive necessity for growth generalizes across environments.

**Supplementary Methods S3 — High-Confidence Determinant (HH) analysis: module construction and dependence-aware conditional perturbation**

The High-Confidence Determinant (HH) analysis integrates module-level organization, conditional perturbation, and feature-level redundancy testing to identify proteins whose quantitative variation is non-redundantly required for accurate growth-rate prediction across environments. Whereas the Global Determinant Analysis identifies proteins with consistent predictive importance across conditions, the HH analysis refines this set by explicitly accounting for feature correlation, some form of constitutive pathway coupling, and shared regulatory structure. All steps in the HH analysis operate on condition-resolved SHAP importance profiles, not on protein abundance.

**Supplementary Methods S4 —Protein-module construction based on SHAP importance profiles**

To organize the 1,857 quantified proteins into coherent analytical units, we clustered them according to the similarity of their condition-resolved SHAP importance profiles. For each protein, the Monte Carlo SHAP procedure produced a 16-dimensional signature summarizing its mean absolute attribution across all aerobic, anaerobic, and anchor conditions. These signatures were standardized and compared using cosine similarity to form a symmetric feature-by-feature similarity matrix. Affinity propagation clustering (damping = 0.80; preference equal to the median of off-diagonal similarities) was applied without pre-specifying the number of clusters, yielding 55 protein modules. Each protein was assigned a fixed module membership for all downstream analyses. All computations were performed with a fixed random seed (42) to ensure reproducibility.

**Supplementary Methods S5 — Module-level conditional perturbation analysis**

To determine whether each module contributes non-redundantly to predictive performance, we evaluated its effect using a dependence-aware conditional perturbation framework applied across all Monte Carlo SHAP iterations. For a given module $c$, protein abundances $X_c$ were modeled as conditionally dependent on the remaining proteome $Z$ via ridge regression ($\alpha = 1.0$) fitted exclusively on training conditions for that iteration. The fitted conditional mean was used to decompose module abundances into predictable, $\hat{X}_c$ and residual components, $R_c$:

$$\hat{X}_c = \mathbb{E}[X_c \mid Z] \qquad (1)$$

$$R_c = X_c - \hat{X}_c \qquad (2)$$

Within each held-out condition, only the residual component $R_c$was permuted, and perturbed inputs were reconstructed as:

$$X_c^{(\pi)} = \hat{X}_c + \pi(R_c) \qquad (3)$$

The Decline-MLP trained in that iteration was then evaluated on the perturbed test set, and the change in predictive error was recorded as:

$$\Delta\mathrm{RMSE}_c = \mathrm{RMSE}\left(f\left(X_c^{(\pi)}, Z\right)\right) - \mathrm{RMSE}\big(f(X)\big) \qquad (4)$$

Repeating this procedure across all Monte Carlo splits (with $n = 364$ iterations) generated a distribution of $\Delta\mathrm{RMSE}_c$values for each module. For each module, we computed the mean effect $\mu$, standard deviation $\sigma$, and standard error:

$$\mathrm{SE} = \sigma/\sqrt{n} \qquad (5)$$

Statistical significance was assessed using a one-sided Normal test of $H_0: \mu \leq 0$, with Benjamini–Hochberg correction (FDR = 0.05). To exclude trivially small effects, we further required $\mu > 2 \cdot \mathrm{SE}$.

Only a minority of modules (13 of 55) produced a statistically significant increase in prediction error upon conditional perturbation (FDR < 0.05), identifying them as significant predictive modules rather than correlated background structure (Fig. 2C). Because growth-limiting constraints may be expressed through distinct proteomic configurations depending on substrate chemistry and oxygen availability, we further required that predictive structure be represented across all environmental contexts. Accordingly, from the full set of modules exhibiting positive perturbation effects (ΔRMSE > 0), we selected the top-ranking module for each substrate–oxygen condition. This procedure yielded a set of 13 constitutive predictive modules that collectively span all environments analyzed and served as the input for all subsequent feature-level conditional perturbation analyses (Fig. 2D).

**Supplementary Methods S6 — Feature-level conditional perturbation and redundancy classification (HH, HL, LH, LL)**

After identifying statistically supported modules, we quantified individual protein contributions using a feature-level conditional perturbation framework analogous to the module-level analysis. For each selected module, every constituent protein $x_j$ was modeled as conditionally dependent on the remaining proteome $X_{(-j)}$ using ridge regression fitted on training conditions only and per each Monte-Carlo iteration:

$$x_j = \hat{x}_j(X_{(-j)}) + r_j \qquad (6)$$

where $\hat{x}_j$ denotes the conditional mean and $r_j$ the residual. Within each held-out condition, only the residual component was permuted, yielding perturbed inputs $x_j^{(\pi)}$, while retaining the measured covariance structure of the full proteome:

$$x_j^{(\pi)} = \hat{x}_j + \pi(r_j) \tag{7}$$

The trained Decline-MLP was evaluated on the perturbed test set, and the resulting change in prediction error was computed as:

$$\Delta\mathrm{RMSE}_j = \mathrm{RMSE}(f(x_j^{(\pi)}, X_{-j})) - \mathrm{RMSE}(f(X)) \tag{8}$$

Repeating this procedure across all Monte Carlo iterations generated a distribution of $\Delta\mathrm{RMSE}_j$ values for each protein. For each protein, we computed the Monte Carlo mean effect size $\mu_j$ of $\Delta\mathrm{R\bar{M}SE}_J$, standard deviation $\sigma_j$, and standard error. A one-sided Normal test of $H_0: \mu_j \leq 0$ was performed using the z-statistic $z_j = \mu_j / \mathrm{SE}_j$, and p-values were adjusted using the Benjamini–Hochberg procedure (FDR = 0.05). To ensure that selected proteins exhibited both statistical support and meaningful effect size, we imposed a joint criterion requiring $q_j < 0.05$ and $\mu_j > 2\,\mathrm{SE}_j$. Proteins passing both filters were designated as feature-level contributors within their respective modules.

To differentiate proteins whose contributions were robust to the choice of conditioning set from those whose importance depended on local intra-module structure, we repeated the above analysis using two conditioning definitions: (i) conditioning on all proteins outside the focal module, and (ii) conditioning on all other proteins in the dataset. The resulting pair of effect estimates for each protein was subsequently used to classify features into high–high (HH), high–low (HL), low–high (LH), or low–low (LL) importance quadrants. This dual-conditioning procedure provided a reproducible and model-agnostic measure of feature-level contribution that captures both unique and redundancy-mediated predictive structure within the proteome.
